# Effects of spatial attention on iconic memory are primarily driven by costs rather than benefits

**DOI:** 10.64898/2026.02.09.704794

**Authors:** Paul J.C. Smith, Niko A. Busch

**Affiliations:** Institute of Psychology, University of Münster, Germany; Otto-Creutzfeldt-Center for Cognitive and Behavioral Neuroscience, University of Münster, Germany

**Keywords:** spatial attention, attentional cost, iconic memory, visual persistence, visual short-term memory

## Abstract

The role of spatial attention in iconic memory – an early interface between visual perception and short-term memory – remains poorly understood. Across two experiments run in 2025, we investigated how endogenous spatial attention modulates iconic memory using a partial-report paradigm. In both experiments, pre-cues manipulated the allocation of spatial attention before stimulus onset. Stimulus arrays were then briefly presented to one hemifield and followed by a post-cue that probed iconic memory at varying delays after stimulus offset. In Experiment 1 (N = 47), valid attentional cues improved performance at short delays. Performance was modeled with an exponential decay function to dissociate effects on initial stimulus availability at short SOAs, the rate of iconic decay, and later transfer to working memory. This analysis indicated that valid pre-cues increased initial stimulus availability relative to invalid pre-cues. In Experiment 2 (N = 66), a neutral pre-cue condition was added, and post-cues were presented at three delays (0, 120, and 1240 ms). This revealed that performance differences were driven by attentional costs at invalidly cued locations, with no detectable benefits at validly cued locations relative to neutral cues. Together, these results show that spatial attention modulates the earliest measurable phase of iconic memory by shaping the initial sensory trace. The cost-dominated pattern suggests that attention primarily suppresses information at unattended locations rather than enhancing representations at attended locations. This finding challenges the view of iconic memory as a pre-attentive sensory store and indicates that attentional selection operates earlier than previously assumed.

**Significance Statement:** The influence of attention on iconic memory – a high-capacity, ultra-brief sensory store – remains highly debated. We demonstrate that endogenous spatial attention modulates iconic memory at its earliest measurable stage. Critically, this modulation reflects an attentional cost at unattended locations rather than a benefit at attended locations, suggesting that spatial attention acts through inhibitory suppression of irrelevant sensory input. These findings challenge models proposing that early sensory representation is categorically attention-free and establish a suppressive role for attention in shaping visual short-term memory at its earliest stage.

## Introduction

Spatial attention plays a central role in visual processing, shaping how we select relevant information and ignore irrelevant input (Carrasco, 2011; Chun et al., 2011). Directing attention to a location enhances perceptual performance, whereas withdrawing it leads to reduced performance. These opposite effects are often described as attentional benefits at attended locations and attentional costs at unattended locations (Carrasco, 2011; Chun et al., 2011; Posner, 1980). Spatial attention improves the perception and discrimination of basic visual features such as color, shape, and motion (Carrasco & Barbot, 2019; Carrasco et al., 2000; Montagna et al., 2009). It also supports the encoding and maintenance of information in visual short-term memory (Matsukura et al., 2007; Myers et al., 2017; Nobre et al., 2008; Souza & Oberauer, 2016) and even determines whether stimuli reach visual awareness, as illustrated by inat-tentional blindness (Mack, 2003). Despite these robust effects on perception, short-term memory, and awareness, it remains debated whether spatial attention also influences the transitional stage between perception and short-term memory, particularly iconic memory.

Iconic memory is a high-capacity, short-duration visual memory store that follows visual perception and precedes visual short-term memory (Dick, 1974; Sperling, 1960). Functionally, iconic memory keeps information from a brief glimpse available for selection and report, such as registering details from a short glance at a traffic sign while driving. Iconic memory is typically measured with partial-report paradigms, in which a briefly flashed multi-item array is followed by a post-cue at variable stimulus onset asynchronies (SOAs) indicating which item to report (Sperling, 1960). When the array and post-cue appear simultaneously, report accuracy for the cued item is high, whereas at long SOAs it declines to chance level. Critically, performance remains well above chance for SOAs up to a few hundred milliseconds, demonstrating that stimulus information—the iconic trace—is still available during this interval but decays rapidly over time. A related approach, change detection, probes the availability of the iconic trace by presenting two arrays separated by a short interstimulus interval and asking whether a change occurred (Becker et al., 2000).

Performance in these tasks depends on an internal impulse response triggered by stimulus onset, which outlasts the physical stimulus because of the low-pass properties of the visual system (Di Lollo, 1980; Loftus & Irwin, 1998). Stimulus information remains reportable as long as the internal response stays above a critical threshold. The duration of this availability is determined by two factors: the amplitude of the initial response and the rate at which it decays (see Figure 1). By sampling performance across different SOAs, these components can be estimated with a psychometric function (Lu et al., 2005). Within this framework, spatial attention could plausibly modulate iconic memory by amplifying the initial response, slowing decay, or both. However, despite these theoretical possibilities, it remains debated whether spatial attention has any measurable effect on iconic memory at all.

**Figure 1:**
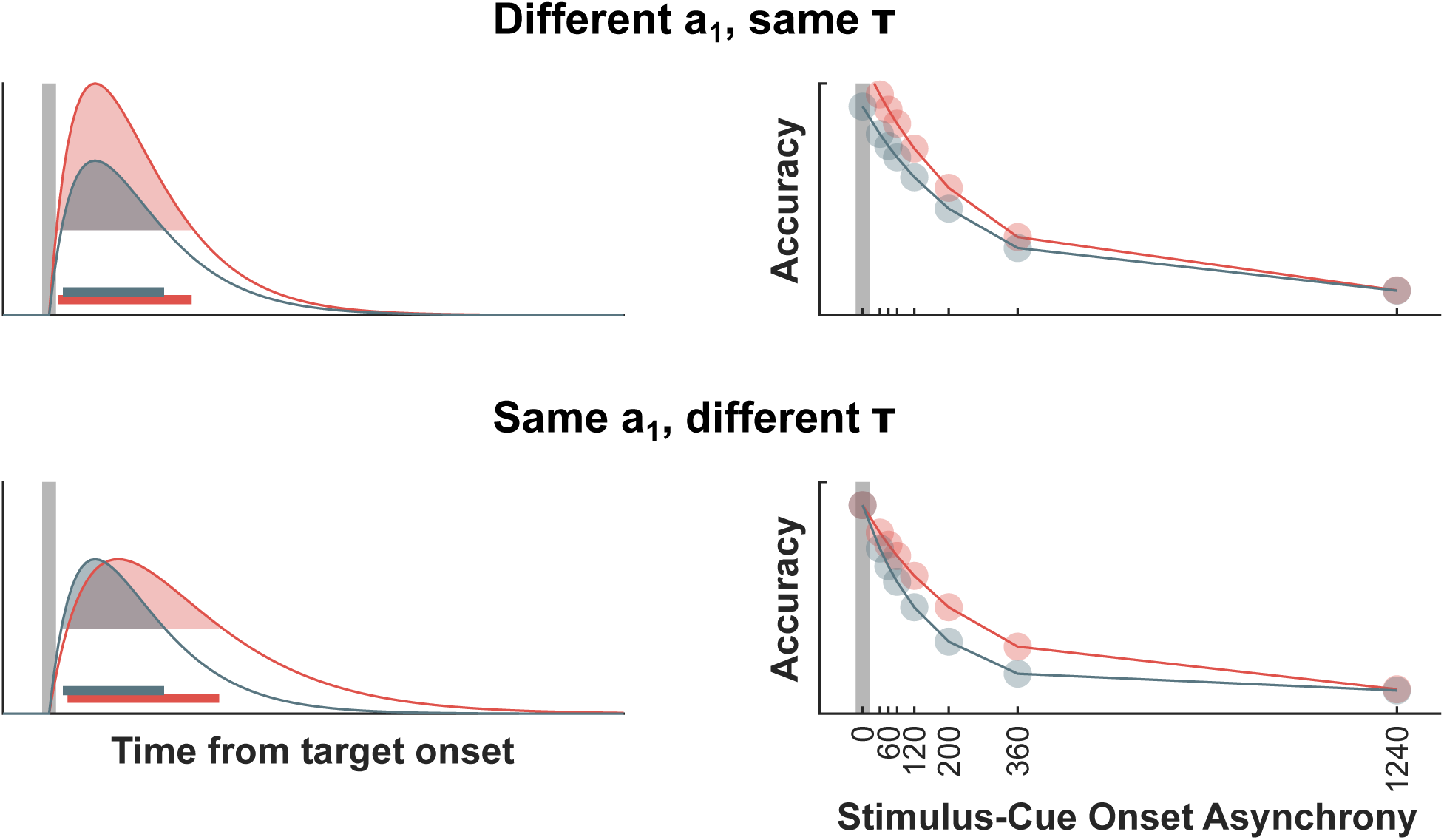
Illustration of two possible mechanisms of temporal persistence modulation. Left: Schematic representation of the impulse response triggered by stimulus onset (indicated by the vertical gray bar). The response shows a rapid onset followed by a gradual decay. The stimulus information persists as long as the response amplitude remains above a critical threshold (shaded area). Horizontal bars indicate the duration of persistence. This persistence can be extended either by increasing the *amplitude* of the initial response (top) or by slowing its *decay* (bottom). Right: Hypothetical performance in the partial report task as a function of stimulus-post-cue SOA. Performance is high at short SOAs and declines with longer SOAs. An amplified response (top) is predicted to enhance initial stimulus availability, thereby boosting performance primarily at short SOAs, which would be captured by the model’s *a*_1_ parameter (see equation 5). A slower decay (bottom) would improve performance at intermediate SOAs, reflected in the model’s *τ* parameter.

A prominent theoretical position arguing that iconic memory is inattentional comes from the distinction between phenomenal consciousness and access consciousness proposed by Block and Lamme. This view holds that iconic memory reflects rich, pre-attentive phenomenal contents (Block, 1995, 2005; Lamme, 2004, 2006). This phenomenal consciousness refers to the subjective, first-person experience of perceptual contents, while access consciousness represents information that is selected and made available for report and subsequent cognitive processing, such as working memory, reasoning, planning, and voluntary control of attention. Within this framework, iconic memory is associated with phenomenal consciousness and is therefore considered to be independent of attentional selection.

A related perspective also assigns a limited role to attention during the earliest stages of visual memory by proposing a fragile visual short-term memory store that follows iconic memory and precedes working memory (Pinto et al., 2017; Pinto et al., 2013; Sligte et al., 2008, 2010). This high-capacity, mask-sensitive store is thought to be more susceptible than iconic memory to attentional interference. Supporting this view, spatial pre-cues in change detection tasks appear to modulate fragile visual short-term memory but not iconic memory itself (Pinto et al., 2017; Pinto et al., 2013). Recent work manipulating feature-based attention in change detection paradigms further suggests that attention affects only fragile visual short-term memory, with no detectable influence on iconic memory (Chiarella et al., 2023; Simione et al., 2019). Together, these accounts maintain that iconic memory is largely immune to the effects of attention, with attention exerting its influence only at later memory stages.

In contrast to these theoretical positions, a growing body of behavioral work indicates that attention can influence iconic memory, although the mechanisms and timing of such effects remain disputed (Chiarella et al., 2023; Gmeindl et al., 2020; Mack et al., 2015, 2016; Persuh et al., 2012; Simione et al., 2019). Persuh and colleagues manipulated attentional load in a dual-task paradigm and found that it reliably affected iconic memory performance. In this paradigm, participants performed an iconic memory task together with a visual search task and were cued only after stimulus presentation which task to perform. High attentional load reduced performance in both partial-report and change detection tasks, suggesting that attention may be required for the formation of iconic memory representations. Increasing attentional diversion by elevating visual-search demands and lowering the probability that the iconic memory task would be cued further degraded performance (Persuh et al., 2012). Mack et al. (2015, 2016) even found inattentional blindness, with observers failing to notice the absence of stimuli in the array when their attention was maximally diverted from the iconic memory task. These inattentional blindness effects have been criticized, however, as possibly reflecting expectation rather than genuine attentional modulation (Aru & Bachmann, 2017; Bachmann & Aru, 2016). Despite this controversy, converging evidence shows that attentional distraction can impair iconic memory performance even when distractors are task-irrelevant and do not act as masks (Botta et al., 2023).

Other studies have examined how the breadth and direction of attention shape iconic memory. Varying pre-cue size in the partial-report paradigm modulates the effective capacity of iconic memory: narrowly focused attention improves performance, whereas a broad attentional focus reduces it (Gmeindl et al., 2020). Spatial pre-cues have also been shown to influence the construction and maintenance of iconic memory representations, with invalid cues inducing clear attentional costs (Botta et al., 2019). However, these results are complicated by a methodological issue: in many paradigms, stimuli are arranged bilaterally around fixation. Valid cues, by restricting attention to one hemifield, reduce the effective memory set size by half compared to neutral cues, which require encoding of the entire array. This set-size confound makes valid trials fundamentally easier than neutral ones, complicating claims about true attentional benefits or costs in iconic memory.

Neural evidence further highlights the complexity of this debate. Electrophysiological investigations of the earliest visual ERP component (C1), often attributed to V1, have traditionally been interpreted as evidence that the first afferent volley of input to V1 is not modulated by attention (Anllo-Vento & Hillyard, 1996; Hillyard & Anllo-Vento, 1998; Morrow et al., 2024), reinforcing the view that attentional influences emerge only after the initial feedforward sweep. However, a recent meta-analysis synthesizing the large and heterogeneous ERP literature concluded that, although individual studies often report null findings, attention overall does exert a moderate overall effect on the C1 (Qin et al., 2022). In contrast, functional magnetic resonance imaging studies have reliably shown attentional modulation of V1 activity (Kanwisher & Wojciulik, 2000). However, such effects may reflect later feedback signals from higher visual areas rather than modulation of the initial feedforward sweep (Martinez et al., 1999). Thus, it remains unresolved whether the earliest sensory responses supporting iconic memory are genuinely attention-sensitive. At the same time, because iconic memory occupies a stage between perception and working memory, both of which are robustly influenced by attention, attentional modulation of iconic memory remains a theoretically plausible and empirically important hypothesis.

A recent study investigating the influence of pre-stimulus brain state on iconic memory provided evidence consistent with a role of spatial attention. Using a partial-report paradigm with variable display-cue SOAs, Smith and Busch (2025b) analyzed ongoing pre-stimulus alpha-band activity (approximately 8-13 Hz), which has been associated with reduced neuronal excitability and corresponding effects on visual perception (Buzsaki & Draguhn, 2004; Dougherty et al., 2017; Ergenoglu et al., 2004; Haegens et al., 2011; Iemi et al., 2017; Limbach & Corballis, 2017; Samaha et al., 2017; Van Dijk et al., 2008). They observed a lateralized effect such that stronger pre-stimulus alpha power over the hemisphere ipsilateral to the to-be-reported stimulus—indicating stronger inhibition—was associated with better performance. This effect was strongest at short display-cue SOAs, indicating a modulation of initial stimulus availability.

Comparable patterns of alpha-band lateralization have frequently been reported in studies using endogenous spatial cueing (Thut et al., 2006; Worden et al., 2000) and are generally interpreted as reflecting inhibition of cortical regions representing task-irrelevant locations. Importantly, because Smith and Busch (2025b) did not employ any pre-stimulus attentional cues, the observed alpha lateralization likely reflected trial-by-trial fluctuations in self-initiated endogenous attention (Balestrieri & Busch, 2022; Bengson et al., 2014; Nadra et al., 2023). Converging evidence comes from pupil-linked arousal, which covaries with attentional state (Alnæs et al., 2014; Mathôt, 2018; Mathôt & Ivanov, 2019) and has likewise been shown to influence initial stimulus availability in iconic memory (Smith & Busch, 2025a). Together, these findings indicate that spontaneous attentional states, indexed by both lateralized alpha power and pupil-linked arousal, modulate the earliest stage of iconic memory.

These results based on spontaneous pre-stimulus brain states lead to several predictions about the effect of spatial attention on iconic memory. First, if spontaneous fluctuations in attention influence iconic memory in the absence of explicit attentional cues, then directing spatial attention with endogenous pre-cues prior to display presentation should produce a comparable effect. Second, attentional cueing should specifically modulate performance at short SOAs between display and partial-report-cue, corresponding to the time window of initial stimulus availability. Finally, this attentional effect should comprise attentional costs for stimuli at unattended locations, consistent with neuronal inhibition of the hemisphere representing task-irrelevant locations. Testing these predictions is particularly important given the ongoing debate about attentional influences on iconic memory, with prior work suggesting that attention exerts its strongest effects at later stages of fragile visual short-term memory and working memory, rather than on iconic memory itself (Botta et al., 2019; Chiarella et al., 2023; Sligte et al., 2008, 2010).

To further clarify the role of spatial attention in iconic memory, we combined a classic partial-report paradigm with attentional cueing across two experiments. Critically, the paradigm involved two distinct cues serving different functions: an endogenous spatial pre-cue presented before display onset to manipulate the allocation of attention, and a partial-report post-cue presented after display onset to probe the contents of iconic memory. In the first experiment, each trial began with an endogenous pre-cue indicating the likely hemifield of an upcoming stimulus array. This was followed by a brief display of six stimuli arranged in a half circle to either the left or right of fixation, and subsequently by a partial-report post-cue appearing after a variable display-cue onset asynchrony. The spatial pre-cue was either valid, correctly indicating the stimulus location, or invalid, indicating the opposite hemifield (see Figure 2A). Performance across SOAs was modeled using a non-linear decay function to characterize the temporal dynamics of iconic memory and to dissociate attentional effects on initial stimulus availability from effects on decay rate (Lu et al., 2005; Smith & Busch, 2025a, 2025b). In a second experiment, we introduced an additional neutral pre-cue that provided no spatial information, serving as a distributed-attention baseline to allow a direct comparison of attentional benefits (valid vs. neutral cues) and attentional costs (invalid vs. neutral cues; see Figure 2B). Importantly, the stimulus array was always presented entirely within a single hemifield, ensuring that memory set size was held constant across cueing conditions. Together with the neutral baseline, this design permits a clean separation of attentional costs and benefits, avoiding confounds present in earlier paradigms (Botta et al., 2019). By combining explicit attentional cueing with a time-resolved analysis of partial-report performance, our approach provides a test of whether and how spatial attention modulates the earliest stages of iconic memory.

**Figure 2:**
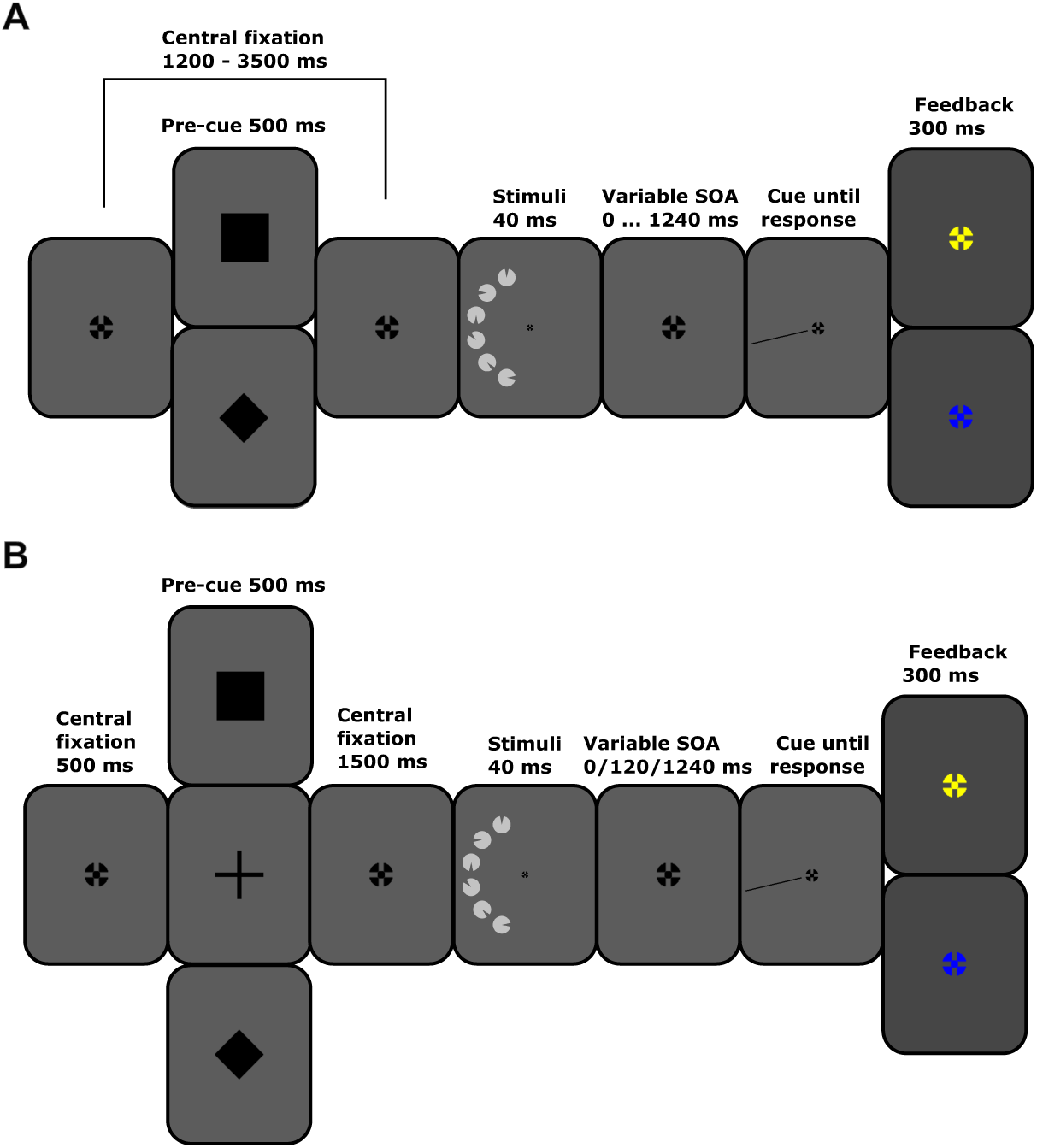
**A**: Illustration of the partial-report paradigm of Experiment 1. Each trial began with a central fixation cross displayed for a variable interval of 1200-3500 ms. During this interval, at least 700 ms before stimulus presentation, a pre-cue, which could either be a rectangle (indicating a stimulus presentation on the right side) or a diamond (indicating a stimulus presentation on the left side), was presented for 500ms. The pre-cue could be valid or invalid. A central fixation cross was presented for the rest of the interval. Following this, a stimulus array consisting of six items was presented on one side of the screen for 40 ms. Participants were then cued to report the orientation of one target item, with the post-cue appearing at various SOAs relative to stimulus onset: at stimulus onset, or 40, 60, 80, 120, 200, 360, or 1240 ms after. There was no time limit for reporting or the subsequent confidence rating. After the confidence rating, feedback was given via the color of the fixation cross, turning blue for a correct response and yellow for an incorrect one. Image proportions are adjusted for illustration purposes. **B**: Illustration of the partial-report paradigm of Experiment 2. Each trial began with a central fixation cross displayed for 500 ms. Following fixation, a pre-cue, which could either be a rectangle (indicating a stimulus presentation on the right side), a cross (indicating no pre-cueing information), or a diamond (indicating a stimulus presentation on the left side) was presented for 500ms. The pre-cue could be valid or invalid. A central fixation cross was again presented for 1500 ms. Following this, a stimulus array consisting of six items was presented on one side of the screen for 40 ms. Participants were then post-cued to report the orientation of one target item, with the post-cue appearing at various SOAs relative to stimulus onset: at stimulus onset, or 120 or 1240 ms after. There was no time limit for reporting or the subsequent confidence rating. After the confidence rating, feedback was given via the color of the fixation cross, turning blue for a correct response and yellow for an incorrect one. Image proportions are adjusted for illustration purposes.

## Experiment 1

### Material and methods

#### Participants

We collected behavioral, ECG, respiratory, and eye-tracking data from 52 healthy participants (mean age 22.94 ± 3.49 years; 38 female, 14 male). We determined the sample size for this experiment and the subsequent experiment based on the sample size of previous studies (Smith & Busch, 2025a, 2025b). All participants had normal or corrected-to-normal vision, reported no history of neurological, psychiatric, or cardiovascular disorders, provided written informed consent, and received either course credit or monetary compensation. The study was approved by the ethics committee of the University of Münster (ref. 2025-07-PS). Data was collected in 2025.

Two participants terminated the experiment early due to headaches, and data from three additional participants were lost due to technical failures. These participants were excluded from all analyses, resulting in a final sample of 47 participants (mean age 22.91 ± 3.63 years; 35 female, 12 male).

#### Stimuli and procedure

The experiment was presented on a 24-inch VIEWPixx/EEG LCD monitor (VPixx Technologies; 1920 × 1080 pixels; 33.76 × 19.38 cm) with a refresh rate of 120 Hz, a pixel response time of 1 ms, and 95% luminance uniformity. The stimuli in the partial-report task were circular shapes with a wedge cut out at one of eight possible orientations (0°, 45°, 90°, 135°, 180°, 225°, 270°, or 315°) extending from the center of the circle to the edge. Circles had a diameter of 2.6° of visual angle (dva). Circles were light gray (RGB: [180 180 180]), and the wedges were dark gray (RGB: [70 70 70]), matching the background color. A symbolic pre-cue indicated the hemifield to be attended: a black rectangle (RGB: [0 0 0]) for right-cued trials and a black diamond for left-cued trials. Both cues matched the size of the fixation cross (0.6 dva). The post-cue consisted of a black line (1 dva long, 0.1 dva thick) pointing to one of the previous stimulus locations and indicated the stimulus to be reported.

Participants were instructed to maintain fixation and to avoid eye movements and blinks during stimulus presentation. Each trial began with a black fixation cross (0.6 dva) presented at the center of the screen for a variable interval (1200–3500 ms). During this interval, the pre-cue was presented for 500 ms, followed by fixation alone for at least 700 ms. Subsequently, six stimuli were arranged along a semicircle in either the left or right visual field and presented for 40 ms. The post-cue appeared either with stimulus onset or after a stimulus–cue onset asynchrony (SOA) of 40, 60, 80, 120, 200, 360, or 1240 ms. Participants reported the orientation of the wedge in the post-cued stimulus using the number pad (e.g., 8 for 0°, 6 for 90°, 2 for 180°). They then rated their confidence as low, medium, or high by pressing 4, 5, or 6 on the number pad. Responses were self-paced. On valid trials (80%), the post-cue pointed to a stimulus in the pre-cued (attended) hemifield. On invalid trials (20%), the post-cue pointed to a stimulus in the opposite hemifield. Feedback was presented 300 ms after the confidence response: correct responses were indicated by a blue fixation cross (RGB: [0 0 255]), and incorrect responses by a yellow fixation cross (RGB: [255 255 0]). A schematic of the trial sequence is shown in Figure 2A.

Participants completed 200 practice trials without data collection, followed by 1000 experimental trials. Short, self-paced breaks were provided after every 200 trials. SOA, stimulus hemifield, and target orientation were counterbalanced. Of the experimental trials, 800 were valid and 200 were invalid. The experiment was programmed in MATLAB R2022b (MathWorks) using Psychtoolbox-3 (Brainard & Vision, 1997; Kleiner et al., 2007; Pelli, 1997).

#### Data acquisition and pre-processing

The experiment was presented on a 24-inch VIEWPixx/EEG LCD monitor (VPixx Technologies; 1920 × 1080 pixels; 33.76 × 19.38 cm) with a refresh rate of 120 Hz, a pixel response time of 1 ms, and 95% luminance uniformity. Recordings were conducted in a dimly lit, sound-attenuated cabin. Participants’ heads were stabilized using a chinrest, with a viewing distance of approximately 86 cm.

Eye movements were recorded using a desktop-mounted EyeLink 1000+ infrared eye-tracking system (SR Research Ltd.) sampling at 1000 Hz (monocular, dominant eye). In addition, electrocardiogram (ECG) and respiratory signals were collected during the experiment but are not analyzed further in the present study.

All data processing and analysis steps were scripted and run in MATLAB R2023a (MathWorks) using custom scripts. Eye-tracking data were processed using the EEGLAB toolbox (Delorme & Makeig, 2004). Continuous eye-tracking data were epoched from −1500 to 1500 ms relative to stimulus onset. Trials were rejected if eye blinks occurred within a −500 to 500 ms window around stimulus onset or if gaze deviated by more than 3 dva from the fixation cross. On average, 229.12 trials were excluded per participant.

Performance and confidence measures were averaged across trials separately for each stimulus–cue onset asynchrony (SOA) and for valid and invalid pre-cue conditions. Sensitivity (*d^′^*), mean confidence, and meta-*I* were computed for each SOA in both cueing conditions.

#### Meta-I

Meta-*I* was quantified as the empirical mutual information between accuracy and confidence ratings, computed as the sum of joint probabilities multiplied by the logarithm of the ratio of joint to marginal probabilities, and expressed in bits (Dayan, 2023) as follows:

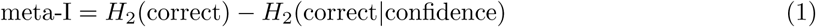

where

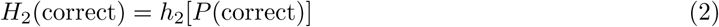

is the entropy of the accuracy,

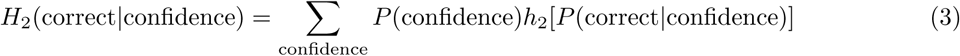

is the conditional entropy of accuracy given confidence;

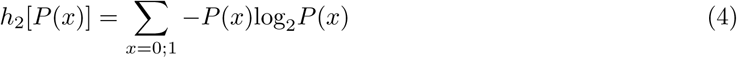

is the entropy of a random variable *x* with two factors e.g., accuracy.

#### Modelling performance

This study tested the hypothesis that endogenous attention modulates the temporal availability of stimulus information in iconic memory. We assumed that the response to a brief visual stimulus follows an impulse response that can outlast the physical stimulus due to low-pass filtering in early visual processing (Di Lollo, 1980; Loftus & Irwin, 1998). As long as the response amplitude remains above a task-specific critical threshold (shaded areas in Figure 1), stimulus information persists and remains available for perceptual and decision-related processes. To assess whether endogenous attention affects stimulus persistence by changing response amplitude or decay rate, we fitted performance data with a nonlinear exponential decay model (Lu et al., 2005):

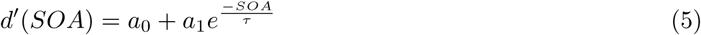

Model parameters were estimated using nonlinear least-squares optimization. The model included three free parameters: *a*_0_, representing sensitivity at long SOAs and reflecting the amount of information transferred into short-term memory in the absence of post-cue benefits; *a*_1_, representing the fast-decaying component of sensitivity associated with initial visual availability; and *τ*, the decay time constant. To obtain stable starting values, the model was first fit to *d^′^* pooled across pre-cue conditions, using initial parameters based on Lu et al. (2005). The resulting parameter estimates were then used as starting values for fitting the model separately to valid and invalid pre-cue conditions.

#### Statistical analysis

Model parameters obtained for valid and invalid pre-cue conditions were compared using Wilcoxon signed-rank tests. Nonparametric tests were chosen because several parameter distributions deviated from normality, as assessed using one-sample Kolmogorov–Smirnov tests.

For analyses of individual SOAs, *d^′^*, confidence, and meta-*I* values were compared between valid and invalid conditions using two-sided paired *t* tests.

#### Transparency and openness

We follow the Transparency and Openness Promotion Guidelines. The data will be made publicly available upon acceptance of the manuscript via the Open Science Framework (osf.io/4b5rt/). Analysis code will be available at github.com/pauljcs/attention-iconic.

## Results

### Analysis of the model parameters

Initial stimulus availability (*a*_1_) was higher for valid than for invalid pre-cues(*z* = 2.38*, p* = 0.0175). The function parameter representing the capacity for information transferred to short-term memory (*a*_0_) was also significantly higher for valid than for invalid pre-cues(*z* = −2.66*, p* = 0.0078). The time constant representing the speed of iconic decay (*τ*) did not show significant differences between valid and invalid pre-cues (*z* = 1.41*, p* = 0.1571) (see Figure 3).

**Figure 3:**
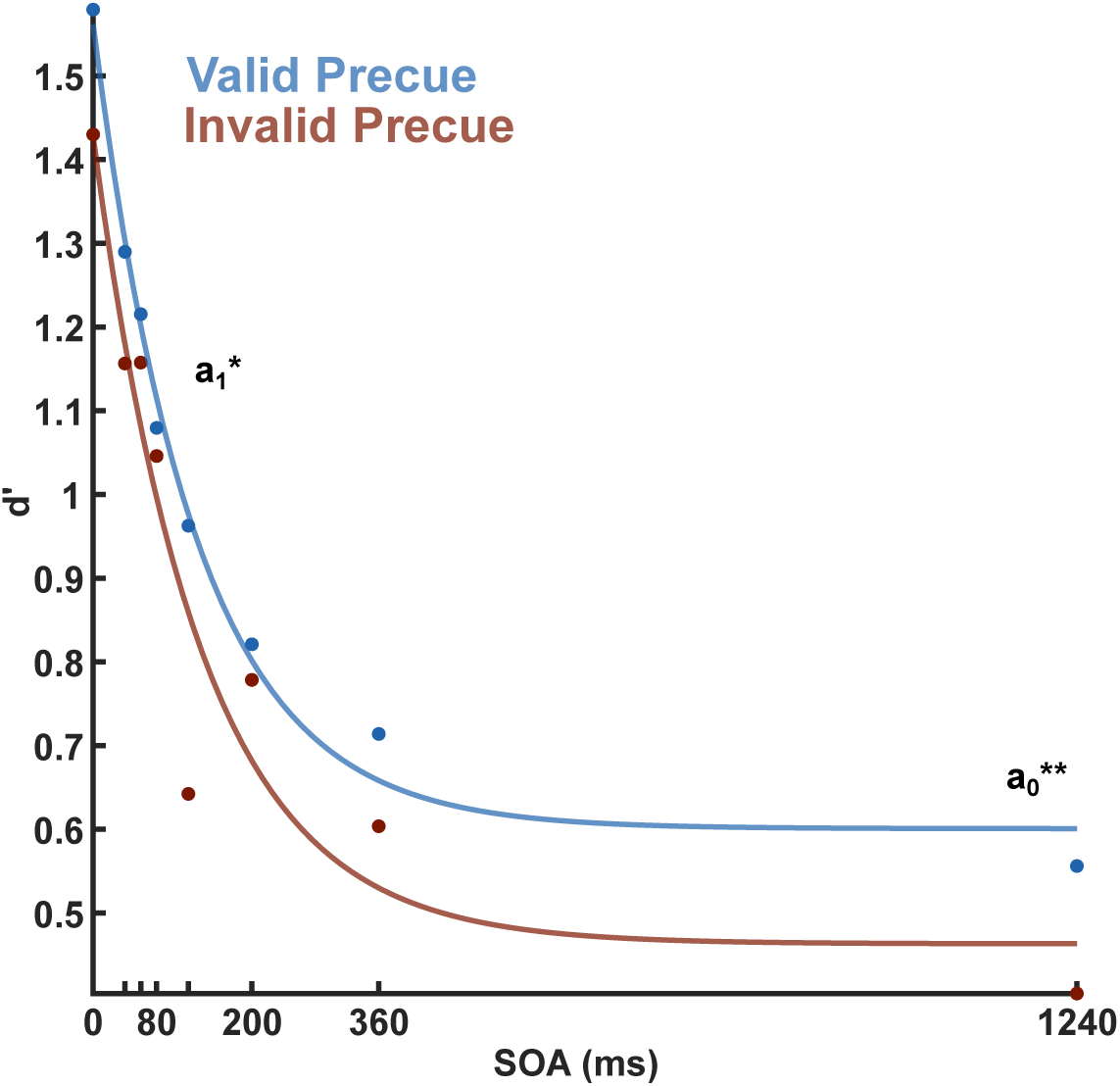
Data from Experiment 1. Accuracy (*d^′^*) data for each SOA. Lines show model fits. Data are shown separately for invalid and valid pre-cues. Model parameters showing a significant difference between the conditions are indicated with asterisks (* = *p <* 0.05; ** = *p <* 0.01).

### Analysis of *d^′^*

The results of the two-sided t-tests for *d^′^* revealed significant differences between invalid and valid pre-cues at SOAs of 0 (*t*(45) = −2.34*, p* = 0.024) and 120 ms (*t*(45) = −3.04*, p* = 0.004), indicating that valid pre-cues led to better performance at these intervals (see Figure 4A and Table 1).

**Figure 4:**
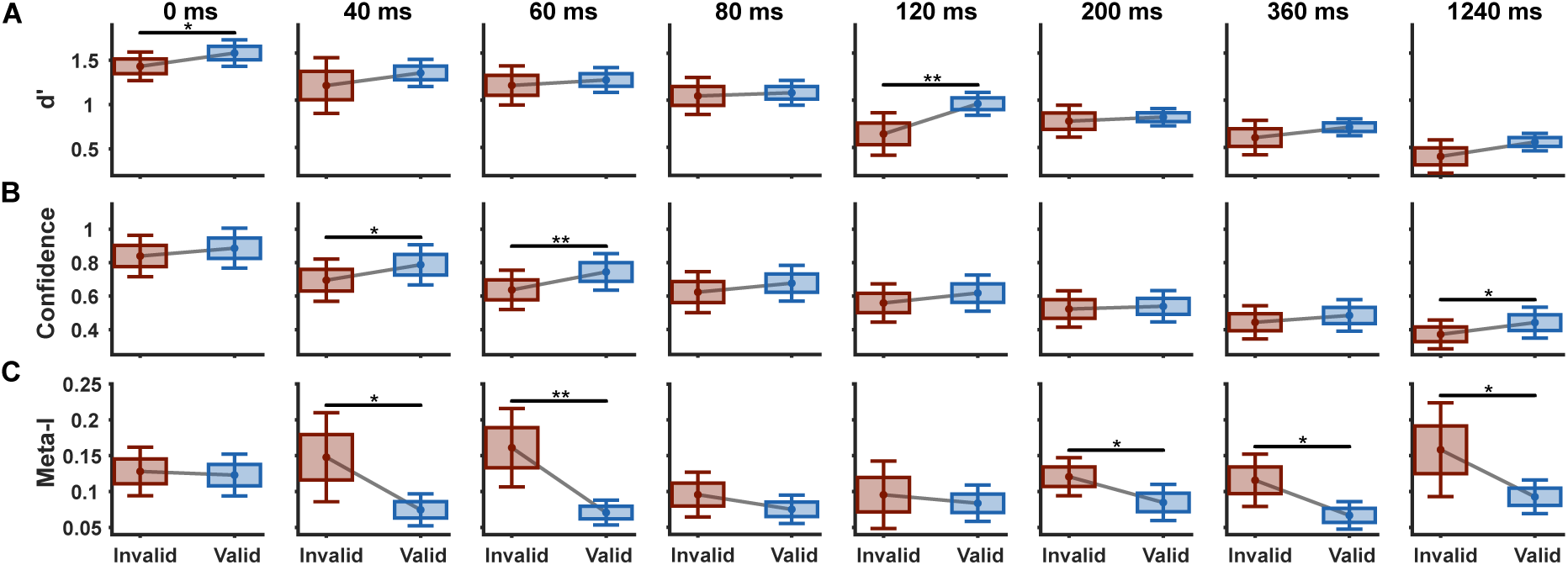
Data from Experiment 1 **A**: Accuracy (*d^′^*) data for each SOA. Data are shown separately for invalid and valid pre-cues. Means of the conditions are connected with black lines. **B**: Confidence data for each SOA. Conventions follow A. **C**: Meta-I data for each SOA. Conventions follow A. SOAs showing a significant difference between the conditions are indicated with asterisks (* = *p <* 0.05; ** = *p <* 0.01).

**Table 1:**
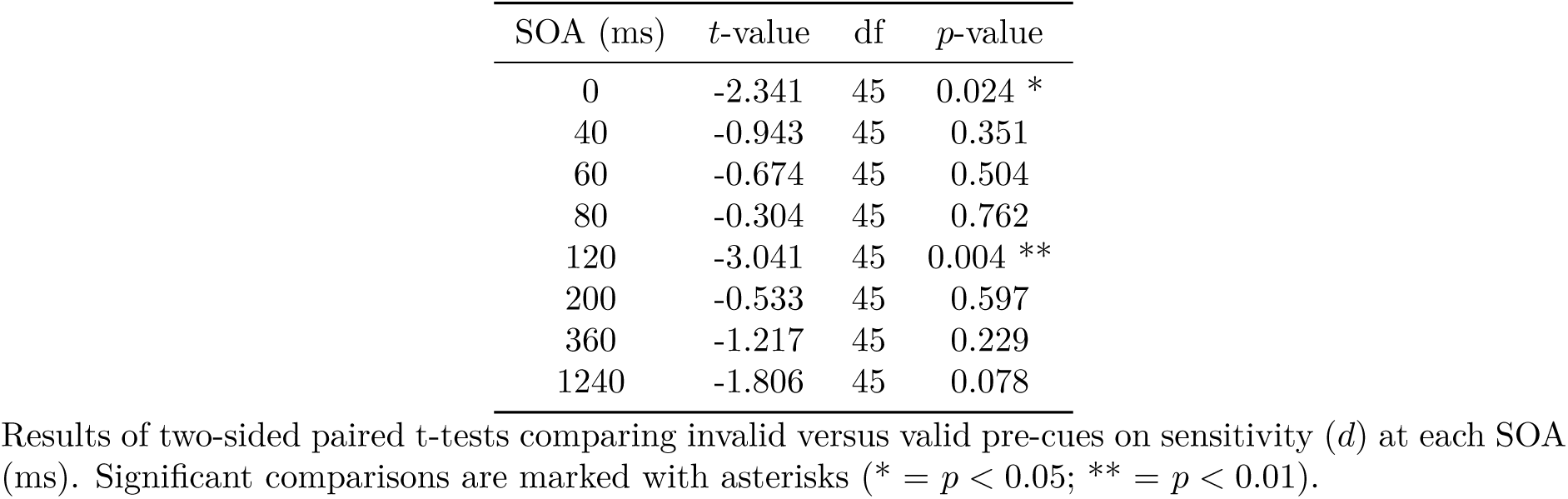
Experiment 1. Results of t-tests for *d^′^* at different SOAs.

### Analysis of confidence

The results of the two-sided t-tests for confidence revealed significant differences between invalid and valid pre-cues at SOAs of 40 (*t*(45) = −2.58*, p* = 0.0129), 60 (*t*(45) = −3.08*, p* = 0.0035), and 1240 ms (*t*(45) = −2.47*, p* = 0.0174), indicating that valid pre-cues led to increased confidence at these intervals (see Figure 4B and Table 2).

**Table 2:** Experiment 1. Results of t-tests for confidence at different SOAs.

| SOA (ms) | $t$-value | df | $p$-value |
| --- | --- | --- | --- |
| 0 | -1.1025 | 45 | 0.2761 |
| 40 | -2.5885 | 45 | 0.0129 * |
| 60 | -3.0819 | 45 | 0.0035 ** |
| 80 | -1.5499 | 45 | 0.1282 |
| 120 | -1.7127 | 45 | 0.0937 |
| 200 | -0.5986 | 45 | 0.5524 |
| 360 | -1.7479 | 45 | 0.0873 |
| 1240 | -2.4703 | 45 | 0.0174 * |

Results of two-sided paired t-tests comparing invalid versus valid pre-cues on confidence at each SOA (ms). Significant comparisons are marked with asterisks (\* = $p < 0.05$ ; \*\* = $p < 0.01$ ).

### Analysis of meta-I

The results of the two-sided t-tests for meta-I between invalid and valid pre-cues showed significant differences at the SOAs of 40 (*t*(45) = 2.44*, p* = 0.0186), 60 (*t*(45) = 3.19*, p* = 0.0025), 200 (*t*(45) = 2.22*, p* = 0.0311), 360 (*t*(45) = 2.48*, p* = 0.0166), and 1240 ms (*t*(45) = 2.10*, p* = 0.0409), indicating that invalid pre-cues led to a significantly better metacognitive evaluation at these intervals (see Figure 4C and Table 3).

**Table 3:** Experiment 1. Results of t-tests for meta-I at different SOAs.

| SOA (ms) | t-value | df | p-value |
| --- | --- | --- | --- |
| 0 | 0.2973 | 45 | 0.7676 |
| 40 | 2.4428 | 45 | 0.0186 * |
| 60 | 3.1979 | 45 | 0.0025 ** |
| 80 | 1.1594 | 45 | 0.2524 |
| 120 | 0.4355 | 45 | 0.6653 |
| 200 | 2.2254 | 45 | 0.0311 * |
| 360 | 2.4874 | 45 | 0.0166 * |
| 1240 | 2.1054 | 45 | 0.0409 * |

Results of two-sided paired t-tests comparing invalid versus valid pre-cues on meta-I at each SOA (ms). Significant comparisons are marked with asterisks (\* = $p < 0.05$ ; \*\* = $p < 0.01$ ).

## Discussion

We observed generally better performance (*d^′^*) for valid than for invalid pre-cues, with significant effects at 0 ms and 120 ms (Figure 4A). Modeling analyses further indicated that valid pre-cues enhanced both initial stimulus availability and information transfer to working memory (Figure 3). The effect on early stimulus availability suggests that spatial attention modulates iconic memory performance at an earlier time window than previously reported. This finding contrasts with prior studies showing only minimal attentional effects at early stages and maximal effects at later stages of visual short-term memory (Botta et al., 2019; Chiarella et al., 2023). The effect on information transfer to working memory is expected and consistent with previous evidence demonstrating robust influences of spatial attention on working memory processes (Awh & Jonides, 2001; Schmidt et al., 2002; Theeuwes et al., 2011).

To determine whether the performance difference between invalid and valid trials reflected an attentional cost, an attentional benefit, or both, we introduced a neutral pre-cue as a distributed-attention baseline in Experiment 2 (see Figure 2B). Based on the effects observed in Experiment 1 on initial stimulus availability, we reduced the set of SOAs to 0, 120, and 1240 ms to maximize statistical power and to accommodate the additional neutral pre-cue condition. Owing to the reduced number of SOAs, Experiment 2 did not include modeling analyses. We expected to replicate the significant performance difference between valid and invalid pre-cues. In addition, we hypothesized a significant performance decrement for invalid relative to neutral pre-cues, consistent with the attentional suppression hypothesis proposed by Smith & Busch 2025b.

## Experiment 2

### Material and methods

#### Participants

We collected behavioral, ECG, respiratory, and eye-tracking data from 83 healthy participants (aged 22.43 ± 2.6 years; 63 female, 20 male). All participants had normal or corrected-to-normal vision, reported no history of neurological, psychiatric, or cardiac disorders, provided written informed consent, and were compensated with course credit or monetary payment. The study was approved by the ethics commission of the University of Münster (ref. 2025-07-PS). Data was collected in 2025.

Three participants were excluded because they experienced headaches and terminated the experiment early, and six additional participants were excluded due to technical failures. Eight participants were excluded because the final number of trials after artifact and trial rejection was fewer than 300 (see Data Analysis). The final sample for the main analysis comprised 66 participants (aged 22.08 ± 2.36 years; 49 female, 17 male).

#### Stimuli and procedure

The stimuli used were identical to those in Experiment 1, with the addition of a cross (RGB: [0 0 0]; with the same dimensions as the other pre-cues) as a neutral pre-cue.

Participants were instructed in the same manner as in Experiment 1. The timing of the paradigm was modified as follows. Participants fixated the center of the screen, where a black fixation cross (RGB: [0 0 0]; 0.6 dva) was presented for 500 ms. The pre-cue was then presented for 500 ms, followed by a 1500 ms presentation of the fixation cross. Subsequently, six stimuli were arranged in a semicircle to the left or right of fixation and presented for 40 ms. The post-cue appeared after a variable SOA, either at stimulus onset or 120 or 1240 ms after stimulus presentation. All other aspects of the paradigm remained unchanged. SOA, stimulus position relative to fixation, and target cut-out orientation were counterbalanced across trials. The experiment comprised 175 invalid, 300 neutral, and 525 valid trials. A schematic of the trial sequence is shown in Figure 2B. Stimulus presentation and response collection were implemented in Matlab 2022b (MathWorks) using Psychtoolbox (Brainard & Vision, 1997; Kleiner et al., 2007; Pelli, 1997).

#### Data acquisition

The data acquisition setup was identical to that used in Experiment 1, except that EEG data were recorded from a subset of 59 participants.

#### Data analysis

All data processing and analysis steps were scripted and run in Matlab2024a (mathworks.com) using custom scripts and the EEGLAB toolbox (Delorme & Makeig, 2004).

The continuous eye-tracking data was epoched from -1500 ms to 1500 ms time-locked to stimulus onset. Trials in which eyeblinks were detected in a range of -500 ms to 500 ms around stimulus onset, or in which participants significantly deviated from the fixation cross (3 dva) were rejected. An average of 230.85 trials were excluded per participant.

Performance and confidence data were averaged across the trials for each SOA in the invalid, neutral, and valid pre-cue conditions. For each SOA in the invalid and valid pre-cue conditions *d^′^*, confidence, and meta-I were computed. Meta-I was again computed as in study one.

A linear mixed-effect model (LME) predicting *d^′^* via pre-cue level was computed for each SOA (*d^′^*(SOA) ∼ pre-cue + (1|participant ID)). LMEs predicting meta-I at each SOA were also computed (*meta* − *I*(SOA) ∼ pre-cue + (1|participant ID)). Post-hoc Wald F-tests were used to further distinguish significant main effects.

#### Transparency and openness

We follow the Transparency and Openness Promotion Guidelines. The data will be made publicly available upon acceptance of the manuscript via the Open Science Framework (osf.io/4b5rt/). Analysis code will be available at github.com/pauljcs/attention-iconic.

## Results

### Analysis of the *d^′^* values

We fitted three linear mixed-effects models to predict *d^′^* as a function of pre-cue level (invalid, neutral, or valid) separately for each SOA (0, 120, and 1240 ms). These analyses revealed a significant main effect of pre-cue level on *d^′^* at the 0 ms SOA (*F* (2, 195) = 4.55*, p* = 0.0117), the 120 ms SOA (*F* (2, 195) = 7.46*, p* = 0.0007), and the 1240 ms SOA (*F* (2, 195) = 3.33*, p* = 0.0380).

Post hoc Wald *F* -tests revealed significant pairwise differences between pre-cue conditions at all SOAs. At 0 ms, performance differed between invalid and neutral trials (*F* (1, 195) = 8.02*, p* = 0.00512) and between invalid and valid trials (*F* (1, 195) = 5.36*, p* = 0.0216), whereas neutral and valid trials did not differ (*F* (1, 195) = 0.266*, p* = 0.607). At 120 ms, invalid trials differed from both neutral (*F* (1, 195) = 13.83*, p* = 0.000261) and valid trials (*F* (1, 195) = 7.66*, p* = 0.00621), with no difference between neutral and valid trials (*F* (1, 195) = 0.906*, p* = 0.342). At 1240 ms, invalid trials differed from valid trials (*F* (1, 195) = 6.08*, p* = 0.0145), whereas neither the invalid–neutral (*F* (1, 195) = 0.333*, p* = 0.564) nor the neutral–valid contrast reached significance (*F* (1, 195) = 3.57*, p* = 0.0605; see Figure 5A).

**Figure 5:**
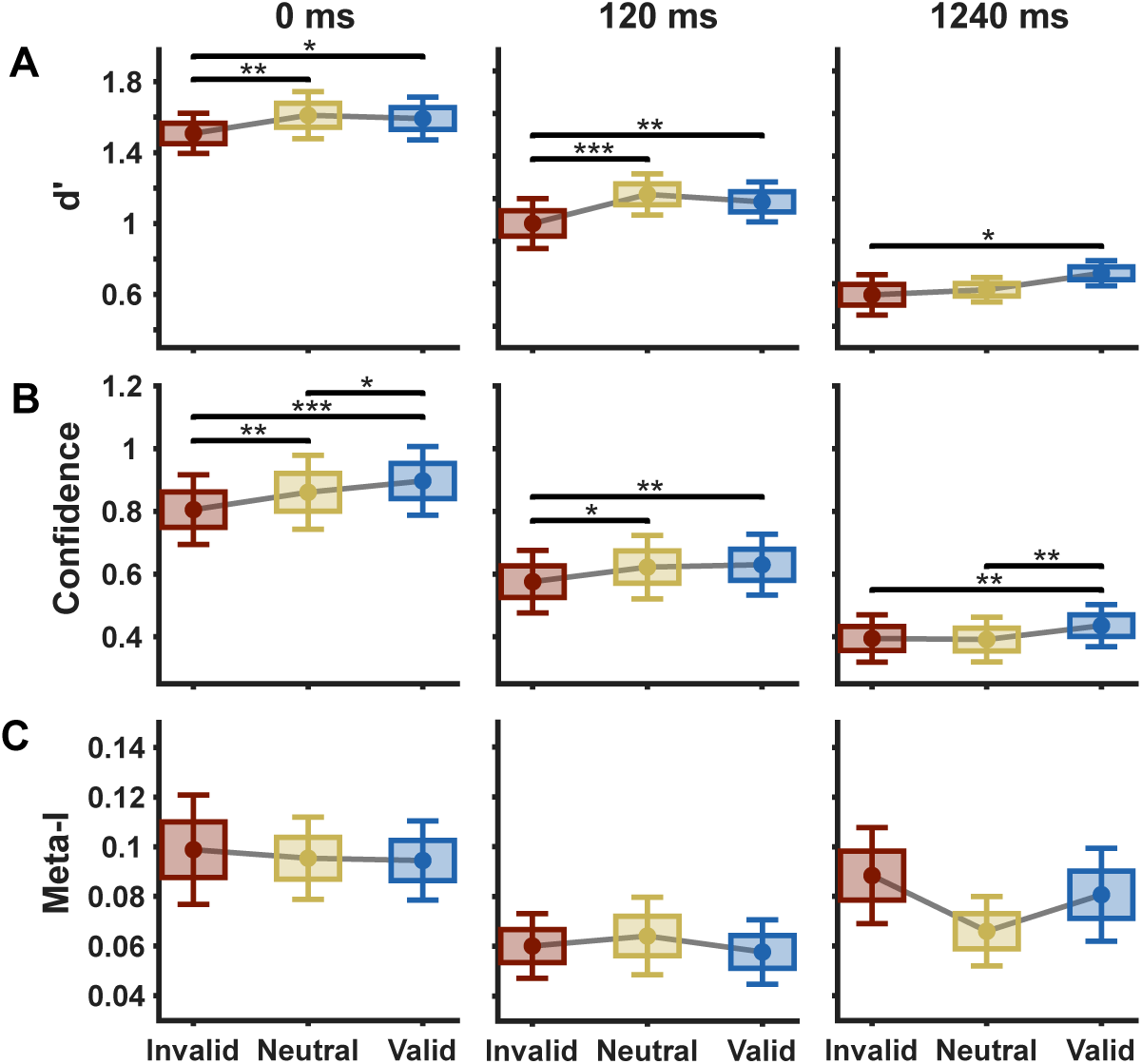
Data from Experiment 2 **A**: Accuracy (*d^′^*) data for each SOA. Data are shown separately for invalid, neutral, and valid pre-cues. Means of the conditions are connected with black lines. **B**: Confidence data for each SOA. Conventions follow A. **C**: Meta-I data for each SOA. Conventions follow A. SOAs showing a significant difference between the conditions are indicated with asterisks (* = *p <* 0.05; ** = *p <* 0.01; *** = *p <* 0.001).

### Analysis of the confidence values

We fitted three linear mixed-effects models to predict confidence ratings as a function of pre-cue level (invalid, neutral, or valid) separately for each SOA (0, 120, and 1240 ms). These analyses revealed significant main effects of pre-cue level at all SOAs (0 ms: *F* (2, 195) = 14.405*, p <* 0.001; 120 ms: *F* (2, 195) = 5.14*, p* = 0.00668; 1240 ms: *F* (2, 195) = 6.04*, p* = 0.00285).

Post hoc Wald *F* -tests indicated that, at 0 ms, all pairwise contrasts were significant, including invalid versus neutral (*F* (1, 195) = 10.416*, p* = 0.00147), invalid versus valid (*F* (1, 195) = 28.386*, p <* 0.001), and neutral versus valid trials (*F* (1, 195) = 4.41*, p* = 0.0370). At 120 ms, confidence differed between invalid and neutral (*F* (1, 195) = 6.46*, p* = 0.0118) and between invalid and valid trials (*F* (1, 195) = 8.78*, p* = 0.00342), whereas neutral and valid trials did not differ (*F* (1, 195) = 0.178*, p* = 0.674). At 1240 ms, the invalid–neutral contrast was not significant (*F* (1, 195) = 0.0580*, p* = 0.810), while both the invalid–valid (*F* (1, 195) = 8.31*, p* = 0.00439) and neutral–valid contrasts (*F* (1, 195) = 9.75*, p* = 0.00206) were significant (see Figure 5B).

### Analysis of the meta-I values

We fitted three linear mixed-effects models to predict meta-I as a function of pre-cue level (invalid, neutral, or valid) separately for each SOA (0, 120, and 1240 ms). These analyses revealed no significant main effect of pre-cue level on meta-I at the 0 ms SOA (*F* (2, 195) = 0.106*, p* = 0.899), the 120 ms SOA (*F* (2, 195) = 0.282*, p* = 0.755), or the 1240 ms SOA (*F* (2, 195) = 2.63*, p* = 0.0743; see Figure 5C).

## Discussion

Experiment 2 replicated the pattern observed in Experiment 1, with invalid pre-cues yielding poorer performance than valid pre-cues across all SOAs. Critically, invalid pre-cues were associated with a significant attentional cost, as evidenced by worse performance relative to neutral pre-cues at the 0 and 120 ms SOAs (see Figure 5A). In contrast, no attentional benefit was observed for valid relative to neutral pre-cues at any SOA. This pattern is consistent with the attentional cost hypothesis proposed by Smith & Busch (2025).

### General Discussion

Does spatial attention influence the earliest stages of visual memory? Although attention has been consistently shown to modulate visual processing, its effects on some early visual processes remain debated (Carrasco, 2011; Kanwisher & Wojciulik, 2000; Qin et al., 2022). In particular, the role of attention in iconic memory has been contentious, with no consensus on whether, how, or when attention influences this earliest stage of visual memory (Botta et al., 2019; Botta et al., 2023; Chiarella et al., 2023; Mack et al., 2015, 2016; Persuh et al., 2012; Pinto et al., 2017; Quilty-Dunn, 2020; Simione et al., 2019). In the current study, we contribute to this ongoing debate by showing that spatial attention modulates iconic memory earlier than previously thought. Based on findings that spontaneous pre-stimulus neural states—classically associated with attentional allocation—predict subsequent iconic memory performance, we hypothesized that spatial attention selectively influences the initial availability of stimulus information, modulates performance specifically at short display–cue SOAs, and incurs a cost at unat-tended locations relative to a neutral baseline (Smith & Busch, 2025a, 2025b).

The introduction of a symbolic attentional pre-cue in the partial-report paradigm reliably altered performance across two experiments (see Figure 4A and Figure 5A). In Experiment 1, valid pre-cues led to better performance at early SOAs than invalid pre-cues (see Figure 4A). Computational modeling revealed enhanced initial stimulus availability for valid relative to invalid pre-cues (see Figure 3). Whereas attentional effects have been demonstrated at later stages of the early visual sensory memory process, especially fragile visual short-term memory (Botta et al., 2019; Chiarella et al., 2023; Pinto et al., 2017; Pinto et al., 2013), our results indicate that spatial attention affects iconic memory at an earlier stage (120 ms SOA) than previously reported (see Figure 3 and Figure 4A). Specifically, our results are consistent with an attentional enhancement of the initial sensory impulse response, rendering stimulus information more readily available for report and thereby improving performance. These findings refine current accounts of attentional modulation in visual memory, as iconic memory – often linked to early visual cortex activity and pre-attentive representations – is often assumed to be largely independent of higher-order processes such as spatial attention (Block, 2011; Teeuwen et al., 2021). In addition, we observed an effect of spatial attention on the parameter indexing transfer of information to working memory, with valid pre-cues enhancing this transfer (see Figure 3). This is consistent with broader evidence that spatial attention improves working memory performance (Awh & Jonides, 2001; Schmidt et al., 2002; Theeuwes et al., 2011). Notably, we found no effect of spatial attention on the decay rate of iconic memory, indicating that attentional benefits do not arise from a slowing of stimulus decay (see Figure 3).

Experiment 2 revealed that the effect of spatial attention took the form of an attentional cost associated with allocating attention to irrelevant locations, resulting in impaired performance for invalid pre-cues relative to the neutral, distributed-attention baseline (see Figure 5A). We observed no evidence for attentional benefits at validly cued locations relative to this neutral baseline (see Figure 5A). These findings suggest that, at the stage of iconic memory, performance may already be close to a ceiling such that additional attentional benefits cannot be expressed behaviorally. Consistent with this interpretation, previous work has shown that attentional benefits emerge progressively over the time course of visual short-term memory, with later stages—such as working memory—being influenced by both attentional costs and benefits (Botta et al., 2019).

Previous research has argued for a pre-attentive iconic memory preceding a more susceptible, fragile visual short-term memory (Chiarella et al., 2023; Pinto et al., 2017; Pinto et al., 2013). However, our results question this distinction: the observation of strong attentional costs at the SOA of 0 and 120 ms shows that attention does modulate stages of the iconic memory process previously thought to be unaffected by attentional effects (see Figure 5A). This effect of spatial attention is especially interesting when compared to other mechanisms of attention, such as feature-based attention, which has been shown to affect fragile visual short-term memory and not iconic memory. It can be argued that the lack of an effect of feature-based attention on iconic memory reflects the relatively later time stage at which feature-based attention is thought to operate (Carrasco, 2011; Chiarella et al., 2023). As spatial attention impacts iconic memory and seems to operate at relatively earlier time stages, our results indicate a functional dissociation in the timing and type of attention influencing the different stages of visual memory (Chiarella et al., 2023; Simione et al., 2019). It reinforces the idea that iconic memory representations, linked to V1 activity, are affected by early attentional processes such as spatial attention, while fragile visual short-term memory, often associated with activity in higher visual areas like V4, is modulated by later mechanisms such as feature-based attention (Carrasco, 2011; Chiarella et al., 2023; Donovan et al., 2017; Teeuwen et al., 2021). This suggests a cascade of attentional processes, where different attentional mechanisms modulate memory representations in visual short-term memory sequentially as they are processed along the cortical hierarchy, with spatial attention influencing iconic memory before feature-based attention modulates fragile visual short-term memory (Chiarella et al., 2023; Simione et al., 2019).

Our findings further contribute to the ongoing debate on the role of attention in iconic memory. Pre-attentive accounts, rooted in the distinction between phenomenal and access consciousness, posit that iconic memory reflects rich phenomenal contents that are largely independent of attentional selection (Block, 1995, 2005; Lamme, 2004, 2006). In contrast, fragile visual short-term memory accounts locate the primary influence of attention at stages subsequent to traditional iconic memory (Pinto et al., 2017; Pinto et al., 2013; Sligte et al., 2008, 2010). By comparison, dual-task and distraction studies have argued that attentional resources are required for the formation of iconic memory representations (Botta et al., 2023; Mack et al., 2015, 2016; Persuh et al., 2012). Our results support an intermediate position between accounts that assume attentional independence of iconic memory and those that posit a necessary role of attention in its formation. Specifically, the immediate cost associated with allocating spatial attention to irrelevant locations indicates that iconic memory is not entirely independent of attentional modulation, but is already influenced by attention at early sensory stages. From this perspective, iconic memory appears to incorporate attentional selection processes that constrain which information subsequently becomes available for access, thereby challenging a strict separation between iconic memory and attentional access assumed in theories distinguishing phenomenal from access consciousness (Block, 1995, 2005; Lamme, 2004, 2006).

The presence of a robust attentional cost in the absence of a corresponding attentional benefit suggests that, within iconic memory, the primary functional role of endogenous spatial attention may be to suppress or filter irrelevant information rather than to enhance signal quality at attended locations. The removal of irrelevant visual input is a central mechanism in classical noise-exclusion models of attention (Carrasco, 2011; Dosher & Lu, 2000a, 2000b). Our findings are consistent with the possibility that similar mechanisms operate at the level of iconic memory. Specifically, while orienting spatial attention toward relevant locations did not measurably enhance performance, orienting attention toward irrelevant locations was associated with impaired performance at the relevant location, consistent with an active suppression or misallocation of attentional filtering.

This asymmetry between attentional costs and benefits is consistent with previous work identifying neural signatures of attentional inhibition in iconic memory (Smith & Busch, 2025a, 2025b). Specifically, increased ipsilateral pre-stimulus alpha power relative to the stimulus location has been associated with enhanced initial stimulus availability and improved iconic memory performance (Smith & Busch, 2025b). Pre-stimulus alpha power, and especially its lateralization, has been widely linked to the allocation of spatial attention (Balestrieri & Busch, 2022; Bengson et al., 2014; Foxe & Snyder, 2011; Thut et al., 2006; Worden et al., 2000), with ipsilateral alpha increases commonly interpreted as reflecting spatially selective inhibition of unattended or irrelevant locations. Within this framework, the attentional cost observed when attention was allocated to irrelevant locations may represent the behavioral consequence of such spatially inhibitory processes: misdirected attentional inhibition suppresses task-relevant sensory input, thereby reducing initial stimulus availability. Together, these findings suggest that inhibitory attentional mechanisms, indexed neurally by pre-stimulus alpha lateralization and behaviorally by performance costs following invalid cues, play a central role in shaping the contents of iconic memory.

### Conclusion

In summary, our two experiments show that spatial attention shapes the earliest measurable time window of iconic memory. Endogenous pre-cues increased the initial availability of sensory information without reliably altering its decay, and performance differences at early SOAs were driven by attentional costs rather than benefits. These results challenge the traditional conceptualization of iconic memory as pre-attentive and indicate a reliable modulation of iconic memory via spatial attention. The cost-dominated signature is consistent with inhibitory selection and noise exclusion accounts and supports a cascade model explaining why spatial orienting can act earlier than feature-based attention, which more strongly impacts fragile visual short-term memory. We establish a novel link between attentional cost, noise reduction, and initial stimulus availability in iconic memory.

## Notes

### Competing Interest Statement

The authors have declared no competing interest.

